# Inferring Gene Regulatory Network Based on scATAC-seq Data with Gene Perturbation

**DOI:** 10.1101/2024.10.10.617724

**Authors:** Wei Shao, Yuti Liu, Shuang Zhang, Shuang Chen, Qiao Liu, Wanwen Zeng

## Abstract

Gene regulatory networks (GRNs) are critical blueprints for understanding gene regulation and the intricate interactions that drive biological processes. Recent advances have highlighted the potential of single-cell ATAC-seq (scATAC-seq) data in GRN inference, offering unprecedented insights into how chromatin accessibility plays an important part in gene regulation. However, existing methods often fall short in providing a quantitative and holistic depiction of regulatory relationships, particularly in capturing the strength, direction, and type of gene regulation simultaneously. In this paper, we present a novel approach that addresses these limitations by leveraging genetically perturbed scATAC-seq data to infer more comprehensive and accurate GRNs. Our method advances the field by integrating pre- and post-perturbation chromatin accessibility data, enabling the construction of GRNs that more accurately reflect the dynamic regulatory landscape. Through rigorous evaluation on seven real datasets, we demonstrate the method’s superior performance in reconstructing GRNs with enhanced precision and interpretability. This work significantly contributes to the field by providing a robust framework for GRN inference, with broad implications for understanding gene regulation in complex biological systems.

## I. Introduction

Gene regulatory networks (GRNs) represent intricate networks of interactions between genes and regulatory elements, playing a pivotal role in understanding the mechanisms underlying various diseases [1]. With the rapid advancement of high-throughput sequencing technologies, numerous methods such as SCENIC [2], GRNBoost2 [3], and DeepSEM [4] have been developed for inferring GRNs using single-cell RNA sequencing (scRNA-seq) data [5]. However, epigenomic data, particularly from single-cell ATAC sequencing (scATAC-seq), offer a more direct and informative perspective for studying gene regulation. Despite this potential, the unique characteristics of scATAC-seq data pose significant challenges for GRN inference due to the much higher dimensionality, making it a more complex task than GRN inference from scRNA-seq data.

Current methods for inferring GRNs from scATAC-seq data remain limited in scope, often failing to capture the full complexity of regulatory relationships. For instance, approaches like DeepTFni [6] construct GRNs as undirected graphs, lacking crucial regulatory directional information, while methods such as SCRIP [7] fail to adequately represent the strength and type of gene regulation. This highlights an urgent need for more comprehensive methods to construct GRNs from scATAC-seq data.

With the advent of gene editing technologies, integrating gene purturbation experiments with single-cell omics data has emerged as a promising research direction. Methods such as CellOracle [8] have improved our understanding of intergenic regulatory mechanisms by analyzing single-cell omics data before and after in silico gene perturbations. Similarly, Perturb-ATAC [9], which combines multiplexed CRISPR perturbations with scATAC-seq, assesses the impact of specific gene perturbations on chromatin accessibility, shedding light on the roles of regulatory elements in gene expression regulation. Together, these technologies have the potential to enhance the depth and breadth of GRN inference by enriching the data with additional layers of information.

In light of these advancements, this paper introduces a novel method for accurately and comprehensively inferring GRNs using genetically perturbed scATAC-seq data. Our approach consists of four key steps: **1) Calculation of Gene Activity Matrices (GAMs):** We begin by computing gene activity matrices from chromatin accessibility data obtained before (GAM_WT_) and after (GAM_KO_) gene perturbations. This step transforms the raw scATAC-seq data into a form that reflects the regulatory activity of genes in both wild-type and perturbed states. **2) Construction of the Base GRN:** Using the GAM_WT_, we construct an initial, or base, GRN. This network serves as a starting point, representing the interactions between genes based on their baseline regulatory activities without any perturbations. **3) Simulation of Gene-Specific Perturbations:** We then apply in silico perturbations to the GAM_WT_, propagating these perturbation effects through the GRN. This simulation process generates a new matrix (GAM_simulated_) that reflects the expected changes in gene activity resulting from specific perturbations, thus modeling how the network responds to these perturbations. **4) Optimization and Refinement of the GRN:** Finally, we iteratively refine the GRN by minimizing the mean square error (MSE) between the simulated gene activity matrix (GAM_simulated_) and the actual post-perturbation matrix (GAM_KO_). Using a gradient descent algorithm, we adjust the edge weights within the GRN to optimize its accuracy, ultimately resulting in a more precise and comprehensive network that captures the strength, direction, and type of gene regulation.

The proposed method was rigorously evaluated on seven publicly available datasets from the GEO database, demonstrating its superior efficacy in GRN inference. Our approach offers several key contributions to the field: **1) Enhanced Precision in GRN Construction:** By integrating genetically perturbed scATAC-seq data, our method more accurately captures the interplay between genes and regulatory elements, reflecting real biological responses to genetic modifications. **2) Comprehensive Representation of Regulatory Relationships:** Unlike existing methods, our approach simultaneously captures the strength, direction, and type of regulatory interactions, offering a detailed understanding of gene regulation through iterative refinement aligned with observed gene activity. **3) Robustness Across Diverse Datasets:** The method’s effectiveness across diverse datasets demonstrates its robustness and adaptability, making it a versatile tool for genetic and epigenomic studies. **4) Advancement of scATAC-seq Data Utilization:** Our method effectively addresses the challenges of scATAC-seq data, highlighting its potential in GRN inference and paving the way for future epigenomic studies. **5) Integration of Gene Perturbation for Improved Inference:** The novel incorporation of gene perturbation data enables dynamic modeling of gene regulation, providing a framework to understand the causal effects of regulatory interactions.

In summary, our method offers a powerful approach for GRN inference, leveraging genetically perturbed scATAC-seq data to construct detailed and interpretable gene regulatory networks with significant implications for both research and clinical applications.

## II. Methods

### A. Datasets

We conducted a comprehensive search of the GEO database using relevant keywords such as scATAC-seq, knockout, and CRISPR, resulting in the identification of 851 experimental projects. Following a thorough manual screening process, we selected four projects, comprising a total of seven perturbed scATAC-seq datasets, to serve as the experimental data for this study. These datasets, which include both mouse and human samples, involve gene knockouts or perturbations and are classified into three distinct categories: single-gene knockout, multi-gene knockout, and Perturb-ATAC. Detailed information regarding these datasets can be found in Table I.

### B. Data Preprocessing

Initially, comprehensive pre-processing of scATAC-seq data was conducted using the R toolkit Signac [14], encompassing four primary steps: sample integration, gene annotation, quality control, and normalization. **1) Sample Integration:** A method of merging overlapping peaks was employed to generate a common peak set for duplicate samples and filter out anomalous peaks. Subsequently, Seurat [15] objects were created for the quantized peaks of each sample, and the duplicate sample objects were merged. **2) Gene Annotation:** Gene annotation information from the Ensembl database was added to the Seurat object, including details such as the location, type, and name of each gene. **3) Quality Control:** To minimize noise and error, we filtered out cells with abnormal total sequencing read counts (top and bottom 1%), high nucleosome signal intensity (top 2%) that impedes chromatin accessibility, and low transcription start site (TSS) enrichment scores (bottom 1%). **4) Normalization:** Based on the TF-IDF method, normalization was performed across cells in the horizontal dimension to correct for differences in sequencing depth and across peaks in the vertical dimension to assign higher weights to rare peaks.

### C. Computing Gene Activity Matrix

Inspired by the use of transcription factor (TF) expression as the motif activity of TF in scBasset [16], we utilized gene activity as the primary indicator to characterize chromatin accessibility and potential functional activity. We quantified the activity of each gene within a cell using Signac based on the total fragment counts at the gene promoter, gene body, and the 2kb upstream region of the gene body. Consequently, we constructed gene activity matrices, GAM_WT_ and GAM_KO_, representing pre- and post-perturbation data in a gene-cell format, respectively.

While GAM_WT_ and GAM_KO_ in gene-cell format share identical row dimensions (denoted as *p* genes), they differ in column dimensions (*n*_1_ and *n*_2_ cells, respectively). To facilitate subsequent comparison, we integrated the two GAMs for cell clustering, converting them into a gene-cluster format with matching dimensions based on the clustering results. The entire process is illustrated in Fig. 1(a).

**Fig 1.**
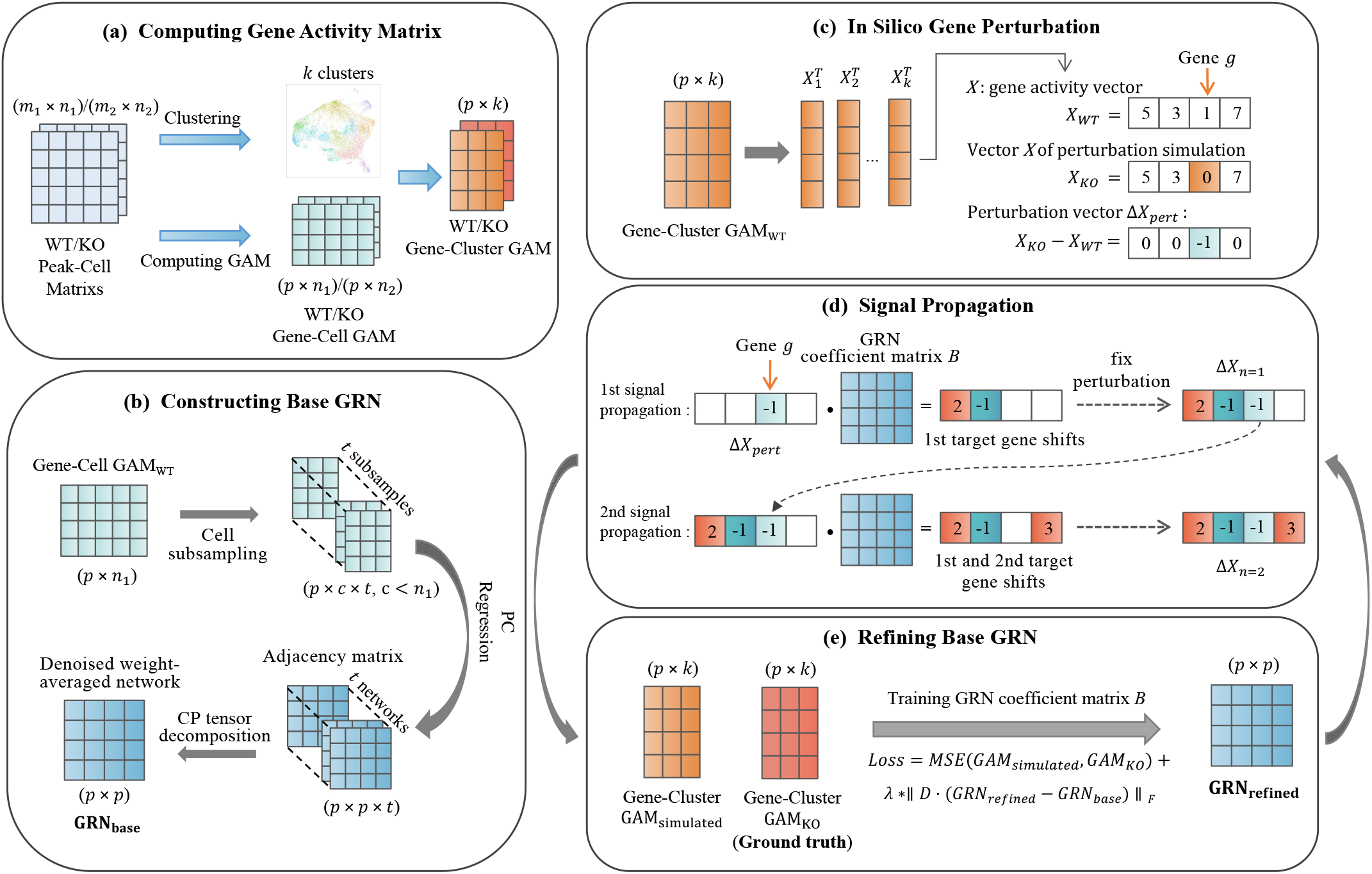
The overall workflow for inferring GRN. (a) Calculate gene activity in the preprocessed data to construct GAMs for pre- and post-perturbation data in gene-cell and gene-cluster formats. (b) Use the wild-type GAM in gene-cell format to infer the base GRN using linear computational methods. (c) In silico perturbation of specific genes in the wild-type GAM in gene-cluster format. (d) Propagate the perturbation effects through the network by multiplying the perturbation change vector with the GRN coefficient matrix. (e) Continuously optimize the GRN coefficient matrix by minimizing the differences between the in silico perturbation and the real perturbation GAMs in gene-cluster format.

To align cellular subpopulations and states shared across sample data, we integrated GAM_WT_ and GAM_KO_ in gene-cell format after batch correction using Seurat [15]. Initially, we conducted feature selection to identify the top 2,000 variable features based on mean-variance calculations by default. Subsequently, these features were scaled, centred, and subjected to PCA dimensionality reduction. Finally, data integration was performed using the CCA algorithm [17].

After data integration, cell clusters were identified using a clustering algorithm based on shared nearest neighbor (SNN) modularity optimization [18]. First, calculate k-nearest neighbors and construct the SNN graph. Then, optimize the modularity function to determine clusters (denoted as *k* clusters). Based on the cell clustering information, the gene activities of cells in each cluster were averaged for both GAM_WT_ and GAM_KO_, converting the GAM to a gene-cluster format with column dimensions of *k*.

### D. Constructing Base GRN

To facilitate the propagation of in silico perturbation effects through the GRN, we use a linear computational approach to construct a directed regulatory network model with quantized weight edges, both positive and negative, as the base GRN. This model’s edge magnitudes indicate regulatory strength, with positive and negative values differentiating regulation types and edge directions representing regulatory flow. We adapted the GRN inference approach from scTenifoldNet [19], using GAM to infer the base GRN after targeted optimizations. The process involves three steps: cellular subsampling, PC regression, and tensor decomposition denoising, as shown in Fig. 1(b).

#### 1) Subsampling of Cells

GAM_WT_ (gene-cell form) contains the activity values of *p* genes in *n* cells. A random subsampling of *c* cells was performed using bootstrapping, and the result was denoted as *S*^*′*^. This subsampling process is repeated *t* times to create *t* subsets of cells, denoted as 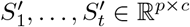, ensuring robustness to cellular heterogeneity within the sample. The values of *t* and *c* are dynamically adjusted based on the size of the dataset, with a minimum requirement that *t × c* exceeds the total number of cells. we set the default values to *t* = 10 and *c* = 500.

#### 2) PC Regression

Each *S*^*′*^ undergoes *p* PC regressions, and the regression coefficients are integrated to form an adjacency matrix *B* with dimensions *p × p*, where the (*i, j*) entry reflects the influence coefficient of the *i*th gene on the *j*th gene. Overall, the risk of overfitting in PC regression is reduced, and computational efficiency is improved by performing regression on *M ≪*PCs (*M ≪ p*, default value of *M* set to 3) to construct the base GRN. The detailed application of PC regression is as follows:

For ease of formula presentation, *S ∈* ℝ ^*p×c*^ is used here to refer to *S*^*′*^ above. The *i*th row of *S*, denoted by *S*_*i*_, represents the activity level of the *i*th gene in the *c* cells. The data matrix *S*_*−i*_ *∈* ℝ ^(*p−*1)*×c*^ is constructed by removing *S*_*i*_ from *S*. First, apply PCA to 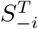 and take the first *M* leading PCs to construct 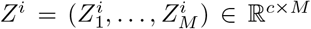, where 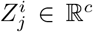 denotes the *j*th PC of 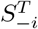 for 1 *≤ j ≤ M*. Mathematically, 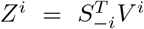, where *V* ^*i*^ *∈* ℝ (*p−*1)*×M* is the PC loading matrix of the first *M* leading PCs, satisfying (*V* ^*i*^)^*T*^ *V* ^*i*^ = *I*_*M*_. Subsequently, the PC regression method regresses *S*_*i*_ on *Z*^*i*^ and solves the following optimization problem:

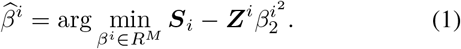

Then, 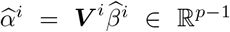, the coefficients of the PC regression model, quantify the influence of the other *p−*1 genes on the *i*th gene. Finally, the coefficients from the *p* regression models are aggregated into a weighted adjacency matrix *B* with dimensions *p × p*. The *i*th row of *B* is 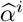, and the diagonal values are all 0. We chose to filter and retain the edges in *B* with the top 90% of absolute magnitude.

#### 3) Denoising via Tensor Decomposition

We stack *t* adjacency matrices to form a third-order tensor Ξ with dimensions *p × p × t*. Applying CANDECOMP/PARAFAC (CP) tensor decomposition, we decompose Ξ into multiple components and use the top components to reconstruct the denoised tensor Ξ_*d*_. Similar to the truncated SVD of a matrix, the CP tensor decomposition assumes that the effective information in Ξ can be described by *d* rank-1 tensors, with the remainder Ξ *−* Ξ_*d*_ primarily representing noise:

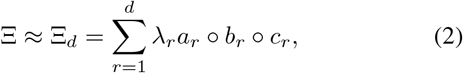

where denotes the outer product, *a*_*r*_ *∈*ℝ^*p*^, *b*_*r*_ *∈*ℝ^*p*^, and *c*_*r*_ *∈*ℝ^*t*^ are unit-norm vectors, and *λ*_*r*_ is a scalar. The reconstructed tensor Ξ_*d*_ *∈*ℝ^(*p×p×t*)^ consists of *t* denoised adjacency matrices, which are averaged to obtain an overall stable adjacency matrix *B*_*d*_. We further normalize entries by dividing them by their maximum absolute value to obtain the final base GRN.

### E. In Silico Gene Perturbation and Signal Propagation

We perform in silico gene perturbations in the gene-cluster form of GAM_WT_ and construct GAM_simulated_ by propagating the perturbation effects step-by-step along the network structure using GRN generated through linear computational methods.

In the gene-cluster form of GAM_WT_, each column of data records the activity values of genes within each cluster. By extracting and transposing each column of data, we obtain the vectors *X*_1_, *X*_2_, …, *X*_*k*_ *∈* ℝ ^1*×p*^. Let *X*_*W T*_ represent the vector corresponding to a cluster before the in silico perturbation. To simulate the knockout of gene *g*, we set the value of gene *g* in *X*_*W T*_ to zero, resulting in the initial simulated perturbation vector, *X*_*KO*_. The process is illustrated in Fig. 1(c).

Let *x*_*i*_ denote the activity value of gene *i* in a particular cluster. Since the base GRN is generated using a linear computational method, if gene *j* is directly regulated by gene *i*, the derivative 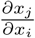 is a constant, denoted as *b*_*i,j*_ in the weight coefficients matrix *B* of the GRN. It represents the weight of the regulatory influence of gene *i* on the target gene *j*:

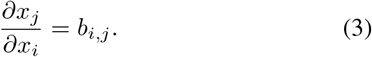

Therefore, the change in the target gene Δ*x*_*j*_ varies with the change in the regulatory gene Δ*x*_*i*_:

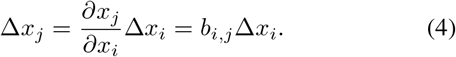

The network edge represents a differentiable linear function, as shown above. The connections between indirectly connected nodes in the network are differentiable composite functions of the linear model. Therefore, we can apply the chain rule to calculate the partial derivatives of the target gene:

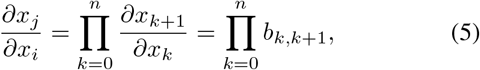

where *x*_*k*_ *∈ { x*_0_, *x*_1_, …, *x*_*n*_*}* denotes the gene activity value of the ordered network nodes on the shortest path from gene *i* to gene *j*. For example, when considering the network edges from gene 0 to 1 to 2, an intermediate connection to gene 1 can be utilized to compute small changes in the response of gene 2 to gene 0:

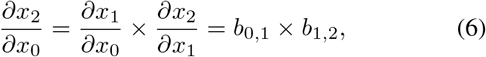

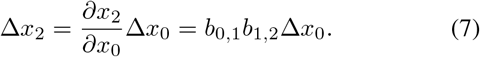

Thus, the change in the activity of the target gene can be calculated as the product of Δ*x*_*i*_ and *b*_*i,j*_. The effect of the in silico perturbation can be propagated linearly by multiplying *X* with *B* in the form:

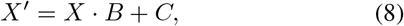

where *X ∈* ℝ^1*×p*^ is a gene activity vector containing *p* genes, and *C ∈*ℝ^1*×p*^ is the intercept vector of the linear model.

The perturbation effect propagation process is illustrated in Fig. 1(d). The product of the activity change vector and *B* is the propagation of perturbation information. Accordingly, by repeatedly performing this operation, the effect of the perturbation on the activity of all downstream target genes after multiple propagations can be deduced. Δ*X*_*pert*_ *∈*ℝ^1*×p*^ is a sparse vector consisting of zeros except for the perturbation target gene *g*:

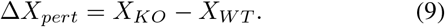

The effect of perturbation on the activity of indirectly regulated target genes from the perturbed target genes to the *n* th layer can be estimated by repeatedly operating *n* times:

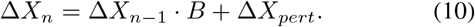

We evaluated seven real datasets to ensure the simulated propagation modeling aligns with real-world perturbations, focusing on the discrepancy between GAM_KO_ and GAM_simulated_ across different propagation rounds. Here we use the untrained base GRN for propagation to better reflect predefined modeling characteristics while excluding the effects of posterior supervised training. Ideally, *n* should balance model complexity and generalization to match biological conditions. For each *n*, we calculated the average Pearson correlation coefficient (PCC) and coefficient of determination *R*^2^ *∈* (*−∞*, 1], with higher values indicating a closer match between the simulated and actual values (Fig. 2, Table II).

**Fig 2.**
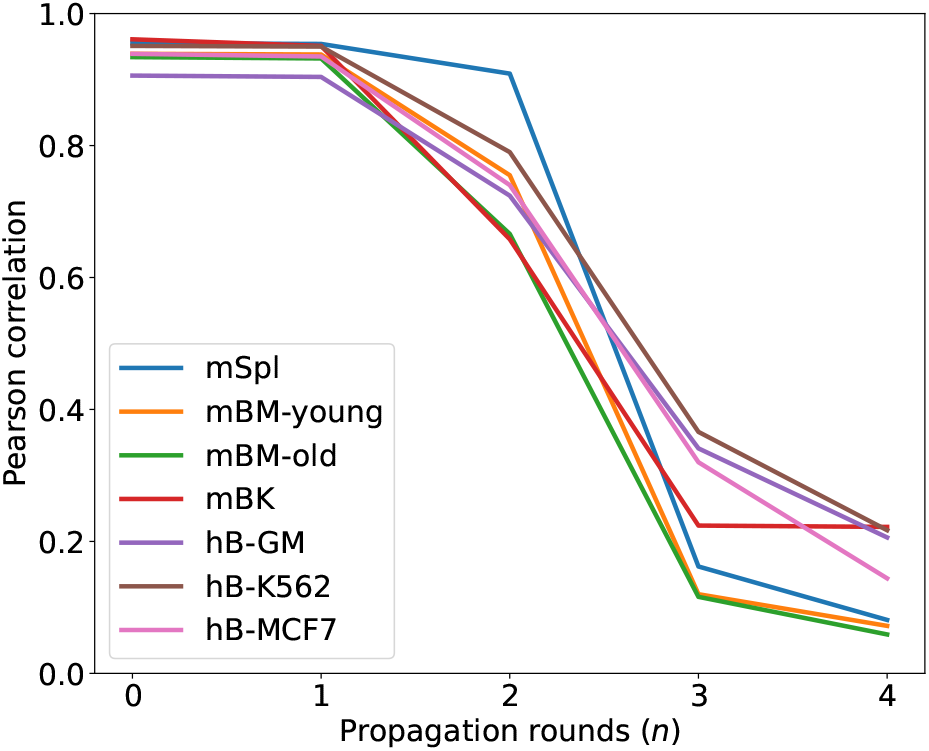
Average Pearson correlation coefficients of the corresponding cluster columns between GAM_simulated_ and GAM_KO_ in the gene-cluster format after in silico perturbation effects have propagated across different propagation rounds *n* on seven real datasets.

**TABLE I.** Description of the datasets.

| Knockout Type | Dataset | Species and Tissues | Cell Type | KO Gene | Cells (WT/KO) | Peaks (WT/KO) | Accession Code |
| --- | --- | --- | --- | --- | --- | --- | --- |
| Single-Gene KO | mSpl | Mouse spleen | CD4+ T - CD8+ T | Prmt5 | 3,506 / 4,475 | 151,569 / 153,810 | GSE195882 [10] |
|  | mBM-young | Mouse bone marrow | HSPC | Tet2 | 10,167 / 5,147 | 179,291 / 162,666 | GSE183675 [11] |
|  | mBM-old |  |  |  | 11,576 / 6,263 | 166,895 / 172,705 |  |
| Multi-Gene KO | mBK | Mouse back skin | HFSC | Foxc1 - Nfatc1 | 7,378 / 8,347 | 228,696 / 232,159 | GSE133646 [12] |
| Perturb-ATAC | hL-GM | Human lymphoid | GM12878 | Multiple genes (40) | 639 / 6,016 | 159,204 / 162,165 | GSE168851 [13] |
|  | hB-K562 | Human blood | K562 |  | 1,187 / 12,354 | 209,024 / 210,069 |  |
|  | hB-MCF7 | Human breast | MCF7 |  | 515 / 4,434 | 251,576 / 259,096 |  |

**TABLE II.** Average *R*^2^ scores of columns between GAM_SIMULATED_ and GAM_KO_ across different propagation rounds.

| Dataset | The $R^2$ scores for n rounds of propagation | | | |
| --- | --- | --- | --- | --- |
|  | n=0 | n=1 | n=2 | n=3 |
| mSpl | 0.498 | 0.500 | 0.399 | **-126.144** |
| mBM-young | 0.433 | 0.420 | 0.055 | **-42.671** |
| mBM-old | 0.008 | 0.028 | -0.722 | **-61.892** |
| mBK | 0.889 | 0.881 | 0.302 | **-148.946** |
| hB-GM | 0.809 | 0.805 | 0.207 | **-472.267** |
| hB-K562 | 0.905 | 0.901 | 0.228 | **-1295.290** |
| hB-MCF7 | 0.876 | 0.866 | 0.071 | **-3890.735** |

**TABLE III.**
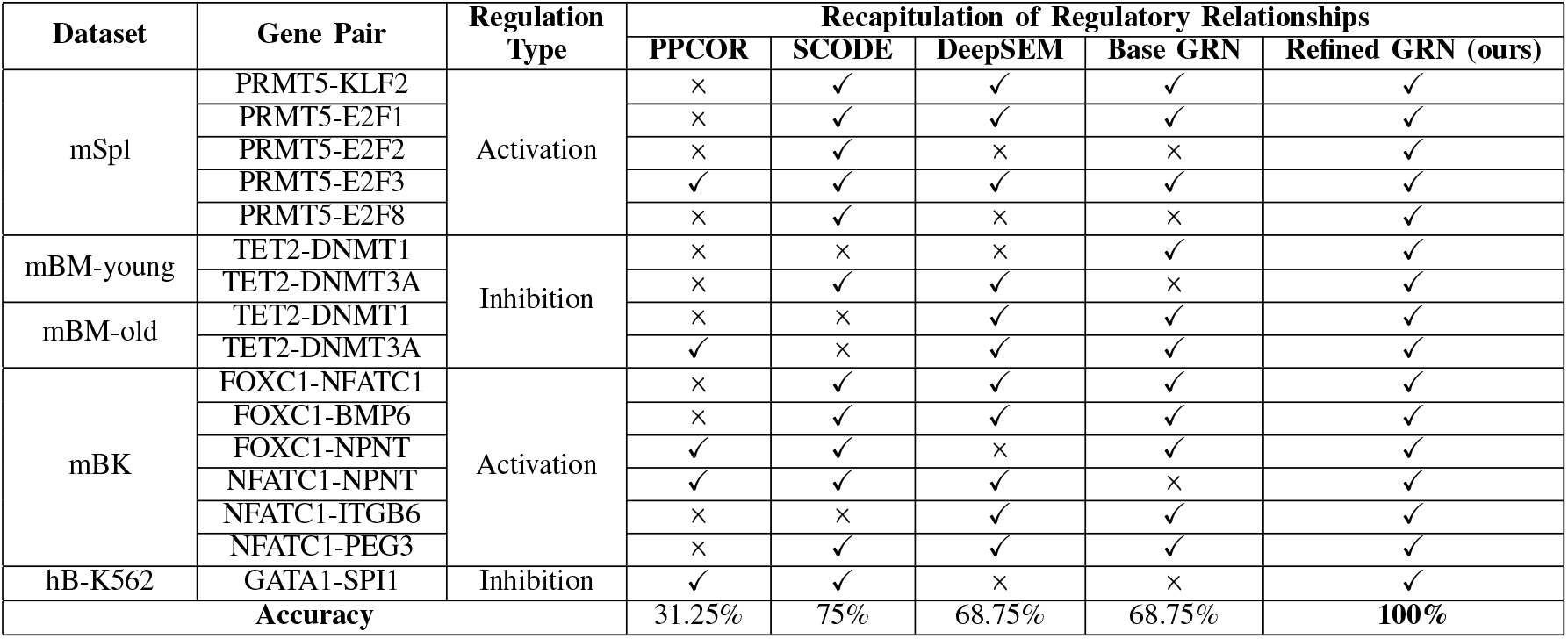
Validation of gene regulatory relationships documented in dataset publications.

In Fig. 2, Round 0 (*n* = 0) compares the wild-type and perturbed GAMs (GAM_WT_ and GAM_KO_), showing only subtle differences and no drastic changes. However, at *n* = 3, both PCC and *R*^2^ values drop significantly into an undesirable range, likely due to the GRN’s tendency to form dense graphs causing rapid accumulation of perturbation effects through feedback loops. While higher *n* improves the model’s fit to regulatory complex relationships, it risks deviating from biological principles. To maintain a reasonable difference between GAM_simulated_ and GAM_KO_, we determined the optimal *n* to be 2.

### F. Refining Base GRN

To refine base GRN constructed by GAM_WT_, we consider the coefficient matrix *B* of the GRN as the trainable parameter matrix for training initialization. The gene-cluster format GAM_simulated_ obtained from in silico perturbations, is compared with the GAM_KO_ (ground truth) corresponding to real perturbations, using the mean square error (MSE) as the discrepancy metric to train and adjust *B*. The Frobenius norm is introduced in the training for regularization constraints to prevent excessive deviation from the base GRN, which could result in a lack of biological plausibility.

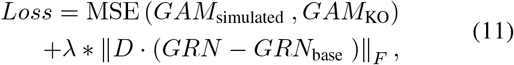

Where *λ* represents the regularization term’s weight, *D* is a diagonal matrix with values of 2 on the diagonal for the corresponding columns and rows of the perturbed genes and values of 1 for the rest of the diagonal entries, assigning greater importance to the direct neighbors of the perturbed genes. The training process employs the ADAM optimizer. The final GRN is the refined GRN obtained by training the base GRN. The workflow is illustrated in Fig. 1(e).

### G. Evaluation Strategies

To evaluate our method’s effectiveness and performance, we compared it against several benchmark methods using seven real perturbed scATAC-seq datasets from the GEO database. The details of datasets are presented in Table I. Given that no existing methods solely use scATAC-seq data to infer GRN with information on regulatory direction, intensity, and type, we converted the scATAC-seq data to GAM. We used mainstream scRNA-seq GRN inference methods as benchmarks. The selected benchmark methods are:

- PPCOR [20]: An R package for calculating partial and semipartial correlation coefficients.
- SCODE [21]: An algorithm for inferring GRN based on ordinary differential equations.
- DeepSEM [4]: A deep generative model that explicitly models regulatory relationships between genes through a neural network version of structural equation modeling.

The primary evaluation criterion is to assess whether the cell type-specific GRNs inferred by each method accurately reconstruct known gene regulatory relationships within the specific cell types. We employed three distinct evaluation approaches in our experiments:

#### 1) Validation Against Dataset Publications

This approach involves assessing whether the inferred GRNs successfully replicate the novel gene regulatory relationships reported in the original research publications associated with the datasets.

#### 2) Cross-Validation Using the TRRUST Database

TR-RUST [22] is a manually curated database containing transcriptional regulatory networks for humans and mice, with 8,444 and 6,552 TF-target relationships for 800 human TFs and 828 mouse TFs, respectively. We evaluated the inferred GRNs by comparing them against the relationships documented in TRRUST using five key metrics: Accuracy, Recall, Precision, F1 score, and Area Under the Precision-Recall Curve (AUPRC).

#### 3) Literature Validation of Refined GRNs

To further validate the refined GRNs, we analyzed the common and unique gene regulatory relationships identified by different methods during the TRRUST evaluation. For relationships uniquely captured by the refined GRNs, we conducted a manual literature review to verify the proportion supported by existing studies, thus reflecting the reliability of the regulatory information provided by the refined GRNs.

## III. RESULTS

To evaluate the ability of our method to reproduce experimentally validated gene regulatory relationships from the original dataset studies, we calculated the prediction accuracy for each method. The comparison results are presented in Table III, where the “base GRN” refers to the initial, unoptimized gene regulatory network generated by our approach, and the “refined GRN” denotes the final, optimized version. It is important to note that the hL-GM and hB-MCF7 datasets were excluded from this evaluation as their corresponding studies did not explicitly validate gene regulatory relationships. Remarkably, our refined GRN demonstrated 100% accuracy, successfully replicating all newly discovered regulatory relationships, representing a 31.25% improvement over the base GRN and a 25% improvement over the best benchmark, SCODE. This significant enhancement not only preliminarily validates the effectiveness of our method but also underscores the potential of the refined GRN in guiding future experimental designs and hypothesis testing.

When scaling the evaluation to encompass 14,996 transcription factor (TF)-target regulatory relationships in the TRRUST database, our method continued to outperform all benchmarks across the five evaluation metrics: Accuracy, Recall, Precision, F1 Score, and Area Under the Precision-Recall Curve (AUPRC). Fig. 3 and 4 illustrate the rigorous results from the TRRUST evaluation. Notably, in test datasets spanning various perturbation categories—including singlegene knockout, multi-gene knockout, and Perturb-ATAC—our refined GRN consistently outperformed all benchmark methods across all metrics, achieving an average improvement of 7.48% in accuracy and 9.30% in AUPRC compared to the best benchmark results. These findings highlight the superior precision and robustness of the refined GRN in inferring gene regulatory networks, reinforcing its effectiveness, accuracy, and adaptability in uncovering genuine regulatory relationships. This comprehensive performance suggests that our approach provides a more reliable foundation for future biological research.

**Fig 3.**
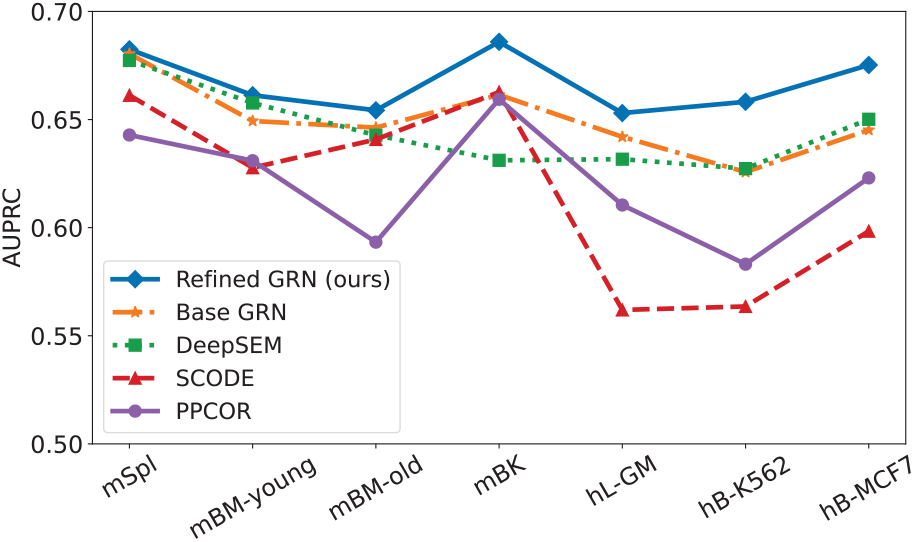
AUPRC results from the TRRUST evaluation experiments.

**Fig 4.**
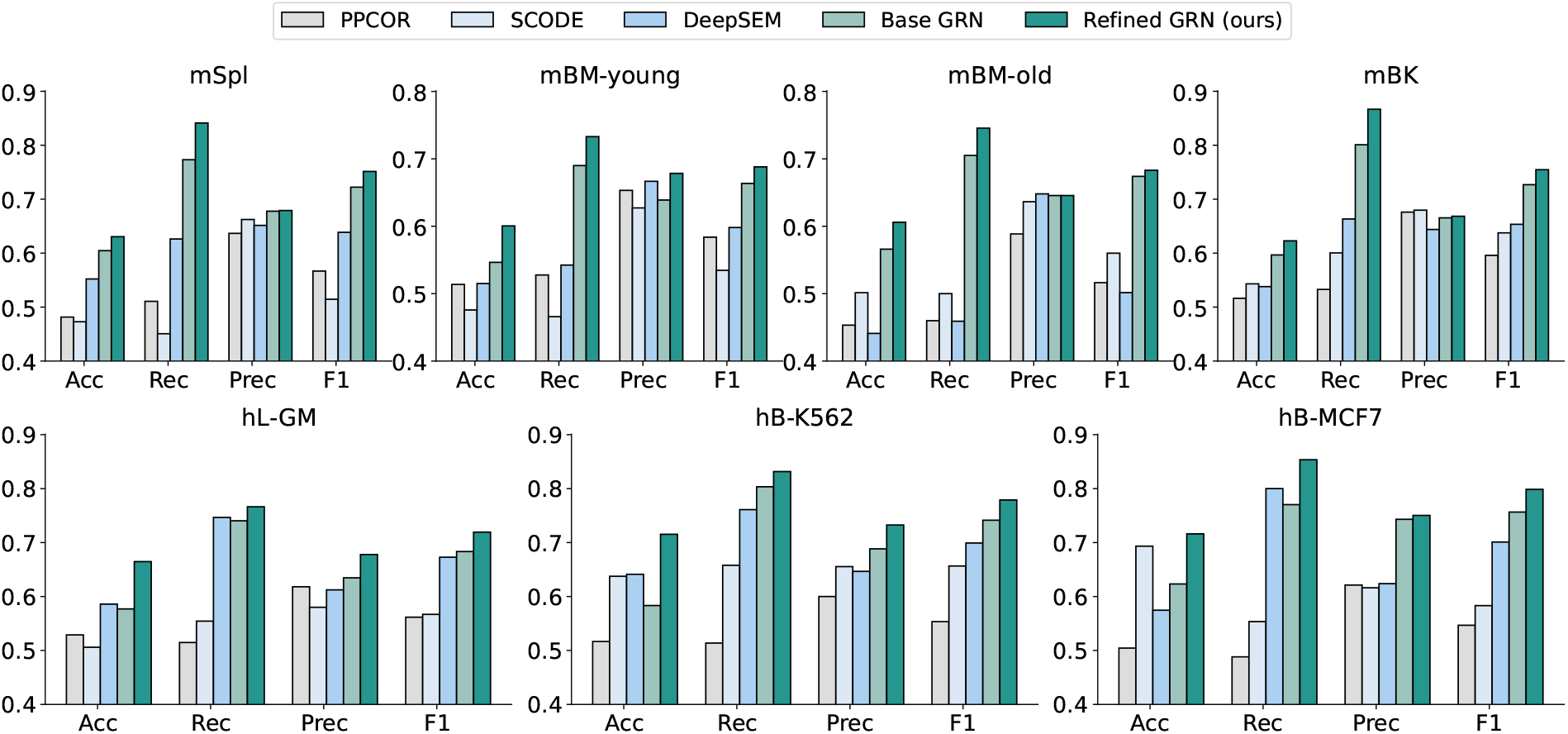
Results from the TRRUST evaluation experiments using seven real datasets. (Acc: Accuracy, Rec: Recall, Prec: Precision, F1: F-score.)

To further explore the common and unique gene regulatory relationships identified by different methods during the TRRUST evaluation, we constructed an UpSet plot (Fig. 5). This plot visually represents the overlap and uniqueness of the inferred regulatory relationships between pairwise comparisons. Each row in the lower part of the figure corresponds to a method, while each column reflects the overlap (or non-overlap) of regulatory relationships between methods. In the datasets analyzed, the refined GRN accounted for 63%, 61%, and 60% of the regulatory relationships in the three respective datasets. The plot clearly shows that the refined GRN not only recapitulates most of the regulatory relationships identified by other benchmark methods but also uniquely captures relationships that other methods fail to identify. This finding further supports the assertion that our refined GRN more accurately and comprehensively reflects authentic gene regulatory interactions.

**Fig 5.**
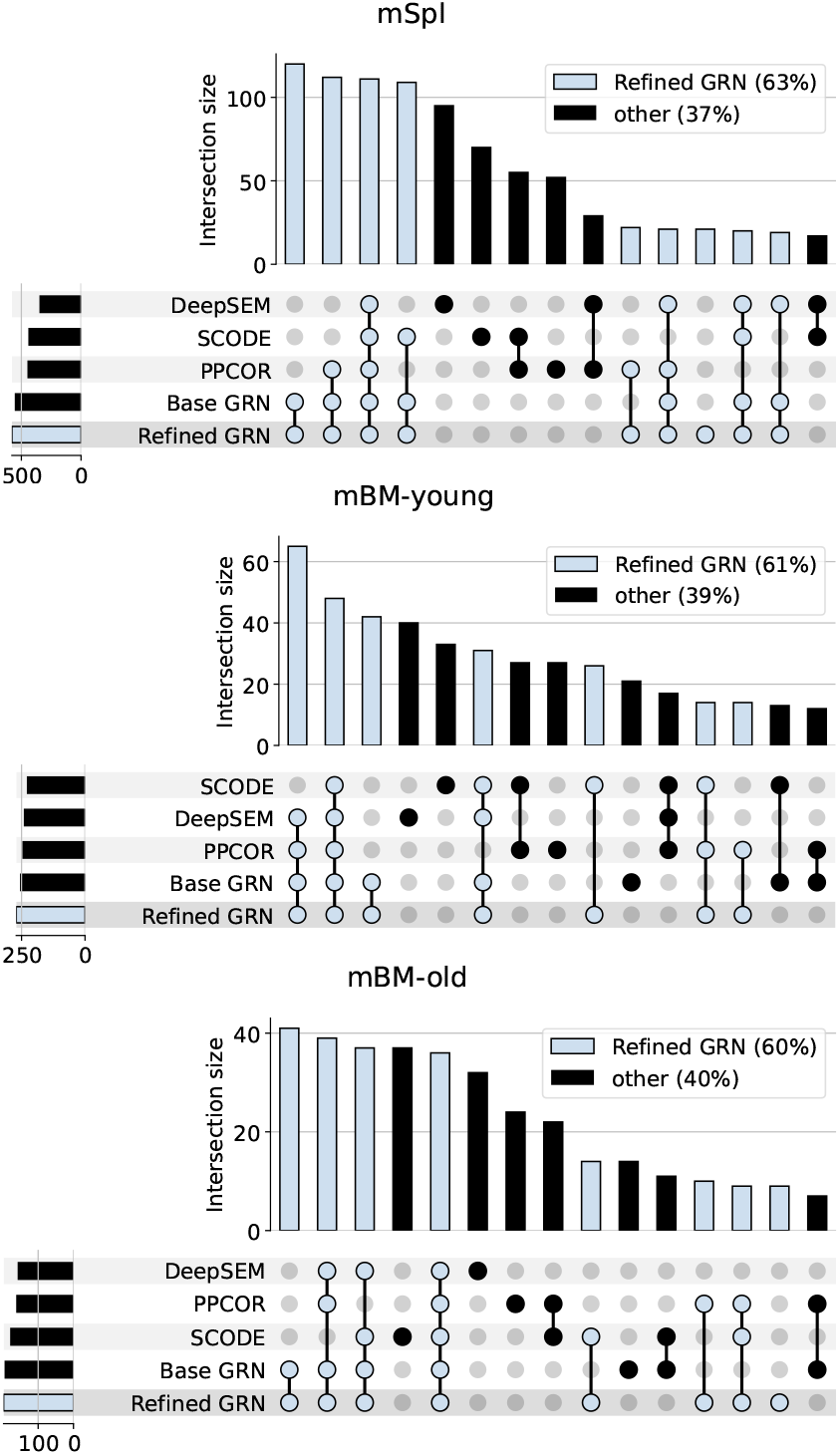
UpSet plots of shared and unique gene regulatory relationships recapitulated in GRNs inferred by different methods in the TRRUST evaluation experiments.

Moreover, we sought to establish the reliability of the regulatory relationships that were uniquely identified by the refined GRN. Given the substantial effort required to manually verify these relationships through literature review, this analysis was conducted on three datasets within the singlegene knockout category. As shown in Fig. 6, a substantial proportion of the uniquely identified regulatory relationships by the refined GRN were experimentally validated in the literature, with 69.23% validation in the mBM-old dataset, 64.71% in the mSpl dataset, and 55.77% in the mBM-young dataset. These results demonstrate that the majority of the gene regulatory relationships uniquely captured by the refined GRN have been previously validated, underscoring the reliability of our approach in identifying accurate, yet potentially undiscovered, regulatory interactions. This provides valuable insights for researchers seeking to explore uncharted gene regulatory landscapes.

**Fig 6.**
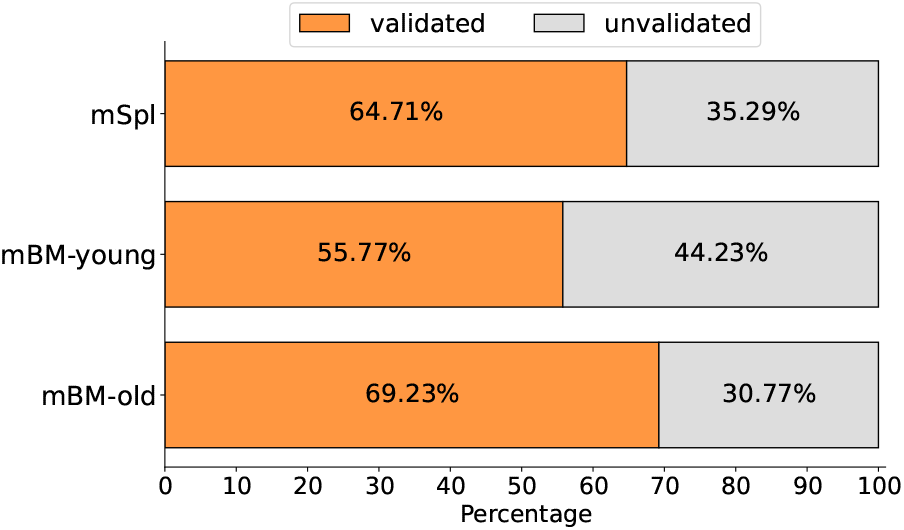
The proportion of gene regulatory relationships uniquely recapitulated by the refined GRN that has been experimentally validated in relevant papers we systematically searched.

In summary, through extensive evaluation and manual validation, we have conclusively demonstrated the effectiveness of our approach in accurately inferring gene regulatory relationships. Our method consistently outperforms existing benchmarks across multiple datasets and metrics, offering a powerful tool for discovering reliable regulatory information that can drive future research and experimental efforts.

## IV. Conclusion

This study presents a novel and robust method for inferring GRNs from genetically perturbed scATAC-seq data, offering significant advancements over previous approaches. Unlike traditional methods, our approach integrates comprehensive quantitative weights alongside detailed regulatory directions and categories, enabling a more precise reconstruction of complex regulatory networks. The workflow, driven by a combination of GAM_WT_ and GAM_KO_, incorporates in silico perturbations and iterative refinement processes to generate an optimized GRN that closely mirrors biological reality. Extensive validation across seven distinct gene knockout datasets demonstrates the superior accuracy, robustness, and adaptability of our method in recapitulating experimentally verified gene regulatory relationships. As technologies like Perturb-ATAC continue to evolve, the availability of genetically perturbed scATAC-seq data will grow, further amplifying the relevance of our approach. This method not only deepens our understanding of intricate gene regulatory mechanisms but also holds immense potential for advancing research in genetics and disease pathogenesis. Our results affirm that this approach sets a new standard for GRN inference, providing a powerful tool for future investigations and practical applications in biological research.

## REFERENCES

[1] G. Karlebach and R. Shamir, “Modelling and analysis of gene regulatory networks,” Nature reviews Molecular cell biology, vol. 9, no. 10, pp. 770–780, 2008.

[2] S. Aibar, C.B. González-Blas, T. Moerman, V. A. Huynh-Thu, H. Imrichova, G. Hulselmans, F. Rambow, J.-C. Marine, P. Geurts, J. Aerts, et al., “Scenic: single-cell regulatory network inference and clustering,” Nature methods, vol. 14, no. 11, pp. 1083–1086, 2017.

[3] T. Moerman, S. Aibar Santos, C. Bravo González-Blas, J. Simm, Y. Moreau, J. Aerts, and S. Aerts, “Grnboost2 and arboreto: efficient and scalable inference of gene regulatory networks,” Bioinformatics, vol. 35, no. 12, pp. 2159–2161, 2019.

[4] H. Shu, J. Zhou, Q. Lian, H. Li, D. Zhao, J. Zeng, and J. Ma, “Modeling gene regulatory networks using neural network architectures,” Nature Computational Science, vol. 1, no. 7, pp. 491–501, 2021.

[5] H. Kim, H. Choi, D. Lee, and J. Kim, “A review on gene regulatory network reconstruction algorithms based on single cell rna sequencing,” Genes & Genomics, vol. 46, no. 1, pp. 1–11, 2024.

[6] H. Li, Y. Sun, H. Hong, X. Huang, H. Tao, Q. Huang, L. Wang, K. Xu, J. Gan, H. Chen, et al., “Inferring transcription factor regulatory networks from single-cell atac-seq data based on graph neural networks,” Nature Machine Intelligence, vol. 4, no. 4, pp. 389–400, 2022.

[7] X. Dong, K. Tang, Y. Xu, H. Wei, T. Han, and C. Wang, “Single-cell gene regulation network inference by large-scale data integration,” Nucleic Acids Research, vol. 50, no. 21, pp. e126–e126, 2022.

[8] K. Kamimoto, B. Stringa, C. M. Hoffmann, K. Jindal, L. Solnica-Krezel, and S. A. Morris, “Dissecting cell identity via network inference and in silico gene perturbation,” Nature, vol. 614, no. 7949, pp. 742–751, 2023.

[9] A. J. Rubin, K. R. Parker, A. T. Satpathy, Y. Qi, B. Wu, A. J. Ong, M. R. Mumbach, A. L. Ji, D. S. Kim, S. W. Cho, et al., “Coupled single-cell crispr screening and epigenomic profiling reveals causal gene regulatory networks,” Cell, vol. 176, no. 1, pp. 361–376, 2019.

[10] Y. Zheng, Z. Chen, B. Zhou, S. Chen, N. Chen, and L. Shen, “Prmt5 deficiency inhibits cd4+ t-cell klf2/s1pr1 expression and ameliorates eae disease,” Journal of Neuroinflammation, vol. 20, no. 1, p. 183, 2023.

[11] T. Hong, J. Li, L. Guo, M. Cavalier, T. Wang, Y. Dou, A. DeLaFuente, S. Fang, A. Guzman, K. Wohlan, et al., “Tet2 modulates spatial relocalization of heterochromatin in aged hematopoietic stem and progenitor cells,” Nature Aging, vol. 3, no. 11, pp. 1387–1400, 2023.

[12] C. Zhang, D. Wang, J. Wang, L. Wang, W. Qiu, T. Kume, R. Dowell, and R. Yi, “Escape of hair follicle stem cells causes stem cell exhaustion during aging,” Nature Aging, vol. 1, no. 10, pp. 889–903, 2021.

[13] S. E. Pierce, J. M. Granja, and W. J. Greenleaf, “High-throughput single-cell chromatin accessibility crispr screens enable unbiased identification of regulatory networks in cancer,” Nature communications, vol. 12, no. 1, p. 2969, 2021.

[14] T. Stuart, A. Srivastava, S. Madad, C. A. Lareau, and R. Satija, “Single-cell chromatin state analysis with signac,” Nature methods, vol. 18, no. 11, pp. 1333–1341, 2021.

[15] Y. Hao, T. Stuart, M. H. Kowalski, S. Choudhary, P. Hoffman, A. Hartman, A. Srivastava, G. Molla, S. Madad, C. Fernandez-Granda, et al., “Dictionary learning for integrative, multimodal and scalable single-cell analysis,” Nature biotechnology, vol. 42, no. 2, pp. 293–304, 2024.

[16] H. Yuan and D. R. Kelley, “scbasset: sequence-based modeling of single-cell atac-seq using convolutional neural networks,” Nature Methods, vol. 19, no. 9, pp. 1088–1096, 2022.

[17] A. Butler, P. Hoffman, P. Smibert, E. Papalexi, and R. Satija, “Integrating single-cell transcriptomic data across different conditions, technologies, and species,” Nature biotechnology, vol. 36, no. 5, pp. 411–420, 2018.

[18] L. Waltman and N. J. Van Eck, “A smart local moving algorithm for large-scale modularity-based community detection,” The European physical journal B, vol. 86, pp. 1–14, 2013.

[19] D. Osorio, Y. Zhong, G. Li, J. Z. Huang, and J. J. Cai, “sctenifoldnet: a machine learning workflow for constructing and comparing transcriptome-wide gene regulatory networks from single-cell data,” Patterns, vol. 1, no. 9, 2020.

[20] S. Kim, “ppcor: an r package for a fast calculation to semi-partial correlation coefficients,” Communications for statistical applications and methods, vol. 22, no. 6, p. 665, 2015.

[21] H. Matsumoto, H. Kiryu, C. Furusawa, M. S. Ko, S. B. Ko, N. Gouda, T. Hayashi, and I. Nikaido, “Scode: an efficient regulatory network inference algorithm from single-cell rna-seq during differentiation,” Bioinformatics, vol. 33, no. 15, pp. 2314–2321, 2017.

[22] H. Han, J.-W. Cho, S. Lee, A. Yun, H. Kim, D. Bae, S. Yang, C. Y. Kim, M. Lee, E. Kim, et al., “Trrust v2: an expanded reference database of human and mouse transcriptional regulatory interactions,” Nucleic acids research, vol. 46, no. D1, pp. D380–D386, 2018.

